# Intratumoral administration of mRNA-1273 vaccine delays melanoma growth in mice

**DOI:** 10.1101/2024.05.06.592840

**Authors:** Dylan T. Boehm, Kaitlyn M. Landreth, Emel Sen Kilic, Katherine S. Lee, Bishal Misra, Sharan Bobbala, F. Heath Damron, Tracy W. Liu

## Abstract

**Background:** Immunotherapies are advantageous for treating cancers; however, their efficacy is limited in unresponsive hosts with “cold” tumor microenvironments. This limitation primarily stems from the lack of infiltrating CD8^+^ T cells, which are major effectors of the anti-cancer immune response. Here, we demonstrate the effects of intratumoral (IT) injections of the COVID-19 mRNA vaccine on the impairment of tumor growth in mice.

**Methods:** Established B16F10 subcutaneous tumor models in wild type mice were used to evaluate the tumor response from IT administration of mRNA-1273. We compared treatment outcomes, immune composition, and transcriptomic effects from mRNA-1273 using survival studies, intravital imaging, flow cytometry, single-cell RNA sequencing, and abscopal studies. The ability of mRNA-1273 to enhance immune checkpoint therapy response was also evaluated.

**Results:** Tumor growth and survival studies following a single IT injection of the COVID-19 mRNA-1273 vaccine showed significant tumor suppression and prolonged survival in tumor-bearing mice. mRNA-1273 treatment resulted in a significant increase in CD8^+^ T cell infiltration into the tumor microenvironment, as observed using intravital imaging and flow cytometry. Further tumor growth suppression was achieved using additional mRNA-1273 treatments. Combination administration of mRNA-1273 with immune checkpoint therapies demonstrated enhanced effects, further delaying tumor growth and improving the survival time of tumor-bearing mice.

**Conclusion:** IT injection of mRNA-1273 significantly reduced tumor growth and enhanced CD8^+^ T cells in the tumor microenvironment. Tumor suppression was further enhanced following multiple injections of mRNA-1273 or when combined with immune checkpoint therapies. This study demonstrates that mRNA vaccines may be used as adjuvants for immunotherapies.

**What is already known on this topic:**

- Most cancer patients develop adequate antibody response to vaccination with mRNA-1273 and produce spike-specific T cell responses. However, it is unclear if the mRNA-1273 vaccine can be used to elicit anti-tumor T cell responses.

**What this study adds:**

- Intratumoral injection of mRNA-1273 significantly delayed melanoma growth.
- Treatment with mRNA-1273 significantly increased CD8^+^ T cell infiltration into the tumor microenvironment.
- Enhanced effects of combination mRNA-1273 and immune checkpoint therapy in anti-tumor immunity.

**How this study might affect research, practice, or policy:**

- mRNA vaccine may be an effective adjuvant with immune checkpoint therapy.

## Introduction

CD8^+^ T cells are the major effector immune cells driving anti-cancer immune responses.^1 2^ Studies correlate more favorable survival in melanoma patients with increased infiltration of CD8^+^ T cells within the tumor microenvironment.^3–5^ The presence of intratumoral T cells are also another major positive predictive factor for immunotherapy response.^6 7^ Tumors with high CD8^+^ T cell infiltration have demonstrated improved responses to immune checkpoint therapy compared to tumors where T cell infiltration is low.^8–13^ Although immune checkpoint therapy has demonstrated a durable response in a subset of melanoma patients, the majority of patients do not respond to immunotherapy.^14–18^ This is thought to occur due to the high prevalence of “cold” tumor microenvironments with low immune T cell infiltration in this type of cancer.^19 20 .^Development of tumor microenvironment modulating strategies that increase CD8^+^ T cell recruitment to the tumor site are needed to convert cold tumor microenvironments to hot, immune cell-rich microenvironments increasing anti-cancer responses.^21 22^

The use of chemokines and cytokines to attract T cells to the tumor site have been explored with varying results.^23–26^ Feasibility of systemic administration of cytokines is limited due to the risk of inducing cytokine storm – a condition where aberrant inflammation can cause damage to host tissues and systems.^27 28^ Alternative indirect routes of boosting cytokine and chemokine responses are an attractive approach due to safety. Vaccines, using low doses of antigens to introduce immunological memory responses without exposure to a whole pathogen, can cause mild and transient increases in cytokines. Thus, interest has risen in the use of intratumoral, rather than systemic, injection of vaccines to enhance T cell recruitment locally to the tumor microenvironment. Several studies utilizing off-the-shelf vaccines have demonstrated promising effects.^29–31^ Intratumoral injection of FDA-approved flu vaccine in combination with tetanus/pertussis/diphtheria vaccines demonstrated tumor regression with increased infiltration of immune cells.^29 30^ There are several reports of clinical regression of tumors following administration of the pertussis acellular vaccine^31^ and the mRNA-1273 COVID-19 vaccine.^32^ In the latter case, intramuscular vaccination with mRNA-1273 resulted in spontaneous tumor regression with localized inflammatory phenotypes in the tumor microenvironment.^32^ This interesting interplay between systemic vaccination and localized effects led to our hypothesis that mRNA-1273 COVID-19 vaccine could drive CD8^+^ T cells into the tumor if administered via an intratumoral injection route. Here, we demonstrate that the intratumoral injection of an FDA-approved lipid nanoparticle-based mRNA-1273 COVID-19 vaccine elicits anti-tumor responses and enhances immunotherapy efficacy. These data pave the path to the next generation of cancer treatments and the development of cancer-related mRNA adjuvant-based therapies.

## Results

### mRNA-1273 immunization results in an increase of CD8^+^ T cells

The onset of the COVID-19 pandemic sparked the widespread use of mRNA-formulated vaccines. SpikeVax (mRNA-1273), has now been extensively studied, and it is well documented that vaccination induces a Th1-skewed immune response.^33 34^ To determine if mRNA-1273 induced a local immune response at the site of administration, we delivered a high-dose intramuscular injection of mRNA-1273 vaccine and observed a significant increase in CD8^+^ T cells in the muscle and lymph nodes 24 h compared to PBS control in healthy C57BL/6 mice (Fig. 1A & B). Given the induced localization of CD8^+^ T cells from mRNA-1273 vaccination, we hypothesized that mRNA-1273 would enhance CD8^+^ T cells into the tumor microenvironment. *In vitro,* we observed that B16F10 melanoma cells express the SARS-CoV-2 spike protein after incubation with mRNA-1273, with detectable protein levels 24 h and 48 h post incubation (Fig. 1C). Uptake of mRNA-1273 by B16F10 cells corresponded to high production of CXCL10, a chemokine involved in T cell recruitment, in culture media (Fig. _S1A)._ ^3^_5-_^3^_8_

**Figure 1.**
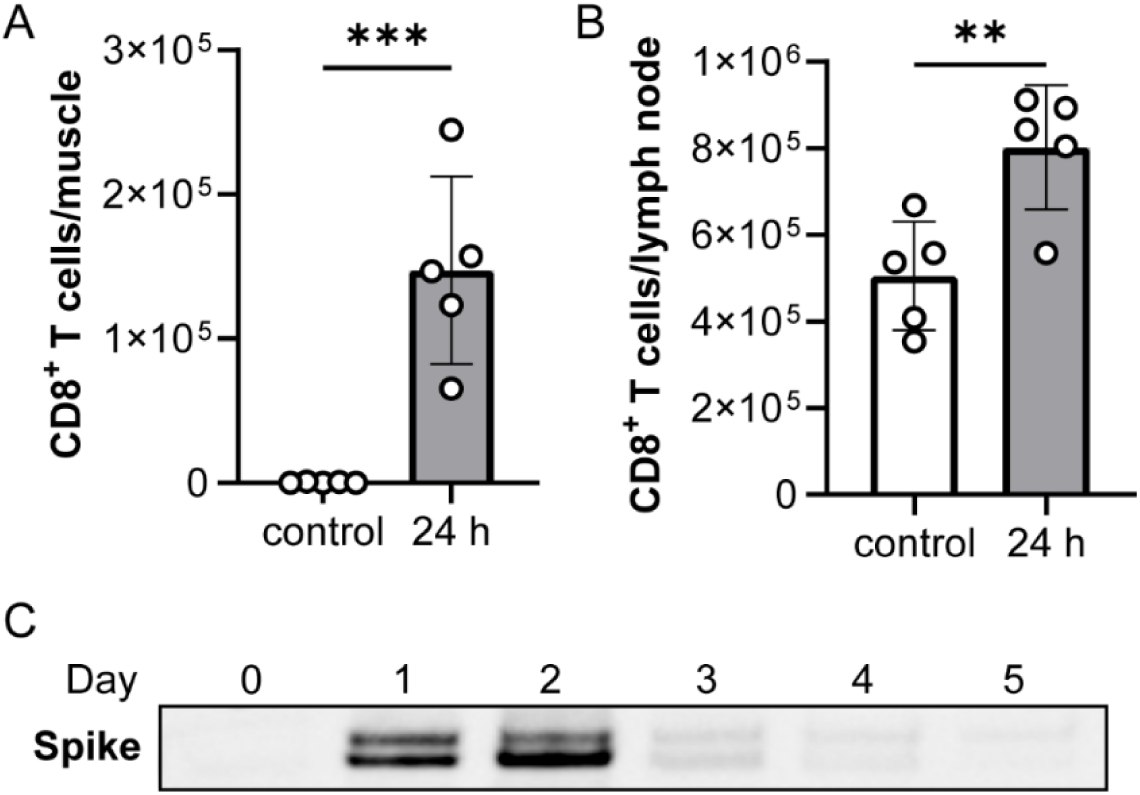
mRNA-1273 vaccine immunization increases CD8+ T cell infiltration at the injection site and is uptaken by B16F10 melanoma cells. Flow cytometry of CD8^+^ T cells in the (A) muscle of injection site and (B) draining lymph node 24 h post immunization with 10 µg of mRNA-1273 or PBS control (n = 5 mice/group). Data shown as mean ± SD; unpaired student t test, **p<0.01, ***p<0.001. (C) Western blot of spike protein in B16F10 cells at different timepoints after incubation with 200 μL (40 μg) mRNA-1273 (n = 3 experiments).

### A single intratumoral injection of mRNA-1273 vaccine significantly delays tumor growth

To evaluate the potential effects of mRNA-1273 vaccination on melanoma tumors, we utilized subcutaneous B16F10 model. When B16F10 tumors reached 5 mm in diameter, an intratumoral injection of 3 μg mRNA-1273 or empty lipid nanoparticle (LNP) without mRNA or PBS occurred. A statistically significant delay in tumor growth was observed in tumors treated with mRNA-1273 compared to those that received LNP or PBS (Fig. 2A – C). While a significant difference in tumor volume was observed between LNP and PBS at 4 days post treatment (Fig. 2B), this difference was lost by day 8 (Fig. 2C). With this delay in tumor growth, we also observed a significant increase in survival time for mice treated with mRNA-1273 compared to PBS (Fig. S1B). To confirm these findings in another melanoma model, we repeated these experiments using the YUMM1.G1 cell line,^39^ and found that IT injection of 3 μg mRNA-1273 again significantly delayed tumor growth compared to PBS (Fig. S1C-E). These data confirm that an IT injection of mRNA-1273 results in tumor reduction.

**Figure 2.**
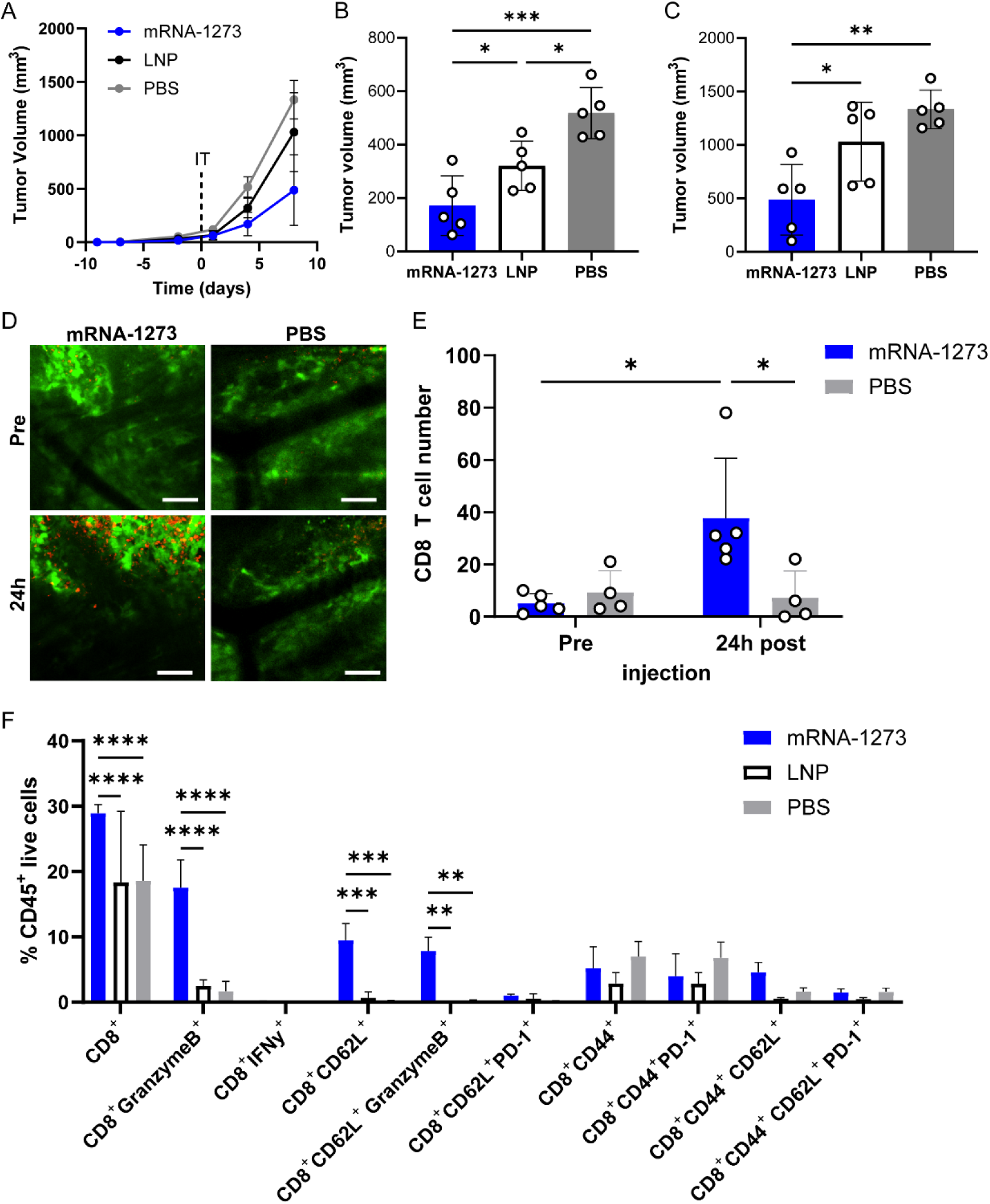
mRNA-1273 vaccine intratumoral treatment reduces tumor growth and increases CD8^+^ T cells in the tumor. (A) B16F10 tumor growth curves and tumor volume measurements at (B) 4 days and (C) 8 days following a 20 µL intratumoral injection of 3 µg mRNA-1273, empty LNP or PBS (n = 5 mice/group). Day 0 represents the time of IT injection when tumors reached 5 mm in diameter. (D) Representative intravital image of tumor (GFP, green) and CD8^+^ T cells (CD8a^+^, red) prior to (Pre) and 24 h post (24h) 10 µL IT injection of 1.5 µg mRNA-1273 or PBS; scale bar represents 100 µm. (E) Corresponding quantification of the number of CD8+ T cells in the tumor (n = 4 – 5 mice per group; n = 3 images per mouse per time point). Flow cytometry analysis of (F) CD8^+^ T cell subsets within the tumor 4 days post IT injection (n = 3 mice/group). Data shown as mean ± SD; one-way or two-way ANOVA followed by Tukey’s multiple comparison test, \**p*<0.05, \*\**p*<0.01, \*\*\**p*<0.001, \*\*\*\**p*<0.0001.

### mRNA-1273 increases CD8^+^ T cell in the tumor microenvironment

Using intravital imaging with skinfold window chamber murine models,^40 41^ we evaluated changes in CD8^+^ T cell frequency within the tumor microenvironment following IT injection of mRNA-1273 or PBS (Fig. 2D). A significant increase in CD8^+^ T cells in B16F10 tumors 24 h post treatment with mRNA-1273 was observed (Fig. 2E). Similar to the subcutaneous tumor studies, a significant reduction in tumor size was observed as well in the intravital studies by quantifying the in B16F10 tumors GFP area, where the fold change normalized to pre-treatment size was < 1 when treated with mRNA-1273, indicating a reduction in tumor size (Fig. S2A & B). Using flow cytometry, the immune cell infiltration profile was evaluated. Similar to the intravital imaging and subcutaneous tumor studies, a significant decrease in B16F10 tumor volume was observed 24 h post IT injection (Fig. S2C). No difference in CD3^+^ or CD11b^+^ cells were observed at 24h following IT injection with mRNA-1273 or PBS (Fig. S2D). We further characterized the immune cell infiltrates in the tumor by flow cytometry at 4 days post IT injection when the largest difference in tumor volume between experimental groups was observed. At 4 days post IT injection, a decrease in CD3^+^ cells (*P* = 0.05) but not in CD11b^+^ cells was observed between mRNA-1273 and LNP (Fig. S2E). It should be noted that the IT injection itself appeared to enhance the infiltration of CD3^+^ or CD11b^+^ cells within the tumor as untreated B16F10 tumors had minimal immune infiltrates - less than 2% of CD45^+^ immune cells were CD3^+^ or CD11b^+^ at the experimental tumor endpoint (Fig. S2F). Interestingly, the majority of the immune cells within the tumor microenvironment at 24 h were CD3^+^CD8^+^ T cells where no differences in population size were observed in tumors treated with mRNA-1273 or PBS (Fig. S2G). Looking at the specific subsets of CD8^+^ T cells, a significant increase in CD8^+^CD44^+^PD-1^+^ T cells was observed in tumors treated with mRNA-1273 (Fig. S2G). However, PBS administration resulted in a significant increase in CD8^+^GranzymeB^+^ T cells compared to mRNA-1273 treatment (Fig. S2G). While the CD4^+^ T cell population made up less than 1% of the immune infiltrate, we observed a significant increase in CD4^+^ T cells in tumors treated with mRNA-1273 (Fig. S2H). A significant increase in CD8^+^ T cells was observed in mRNA-1273 treated B16F10 tumors compared to LNP and PBS controls 4 days post IT injection (Fig. 2F). Approximately two-thirds of the CD8^+^ T cells from mRNA-1273 treatment were Granzyme B^+^, which was a significant increase when compared to LNP or PBS controls (Fig. 2F). Approximately half of these CD8^+^GranzymeB^+^ T cells were CD62L^+^ and was also significantly increased in B16F10 tumors treated with mRNA-1273 compared to LNP or PBS controls (Fig. 2F). All CD8^+^ T cells lacked IFN-γ expression. At 4 days post IT injection, an increase in CD4^+^ T cells was seen in which a significant increase in CD4^+^CD44^+^CD62L^+^ and CD4^+^CD62L^+^ T cells were observed in B16F10 tumors treated with mRNA-1273 compared to LNP or PBS controls (Fig. S2I). The specificity of the CD8^+^ T cells within B16F10 tumors were also evaluated using TRP-2 staining by flow cytometry. A significant increase in TRP-2^+^CD8^+^ T cells was observed at 24 h but was lost by 4 days post IT mRNA-1273 treatment (Fig. S2J & K). Approximately 20% of CD8^+^ T cells were TRP-2^+^ at 24 h post IT injection of mRNA-1273 (Fig. S2J). There were no differences observed in T cell subsets in the spleen at 24 h or 4 days post IT injection of mRNA-1273, LNP or PBS (Fig. S3). No significant differences were seen in T cell subsets in the draining lymph node at 24 h post IT injection of mRNA-1273, PBS or LNP (Fig. S4A & B). Interestingly, there was a significant increase in CD8^+^ T cells, CD4^+^ T cells and CD4^+^CD44^+^ T cells in the draining lymph node following treatment with PBS compared to mRNA-1273 or LNP at 4 days post injection (Fig. S4C & D).

### Multiple mRNA 1273 IT injections lead to enhanced reduction of tumor growth

While a significant decrease in tumor volume following a single mRNA-1273 IT injection was observed, we did not achieve long-term survival or tumor suppression. We evaluated whether greater tumor control or long-term survival could be achieved using multiple mRNA-1273 IT injections. IT injections began when tumors reached 5 mm in diameter and were administered every 3 days. A significant delay in tumor growth was observed following three 20 μL IT injections of 3 μg mRNA-1273 compared to LNP or PBS controls (Fig. 3A). At 8 days post injection, a statistically significant delay in tumor growth was observed in tumors treated with mRNA-1273 compared to LNP or PBS as well as between LNP and PBS (Fig. S5A). A significant increase in survival was observed when B16F10 tumors were treated with three IT doses of mRNA-1273 compared LNP or PBS (Fig. 3B). No differences in survival was observed between treatment with three IT doses of LNP or PBS (Fig. 3B). Compared to a single dose of mRNA-1273, three IT injections significantly decreased tumor volume at day 8 (Fig. 3C). However, no significant increase in survival was observed comparing multiple mRNA-1273 with a single mRNA-1273 dose (Fig. S5B). Whether an IT injection of mRNA-1273 could induce abscopal effects, where treatment of one tumor induces positive effects on a secondary distal tumor, was also evaluated. Mice were implanted with tumors on the right and left flanks, where only the right flank was treated with multiple mRNA-1273 IT injections or PBS control. A significant reduction in both the IT mRNA-1273 treated right and untreated left tumor was observed compared to PBS treated control mice at day 8 post IT treatment (Fig. 3D & S5C). However, the tumor reduction in the abscopal studies was not similarly delayed as that observed when a single tumor was treated with multiple mRNA-1273 doses (Fig. S5D).

**Figure 3.**
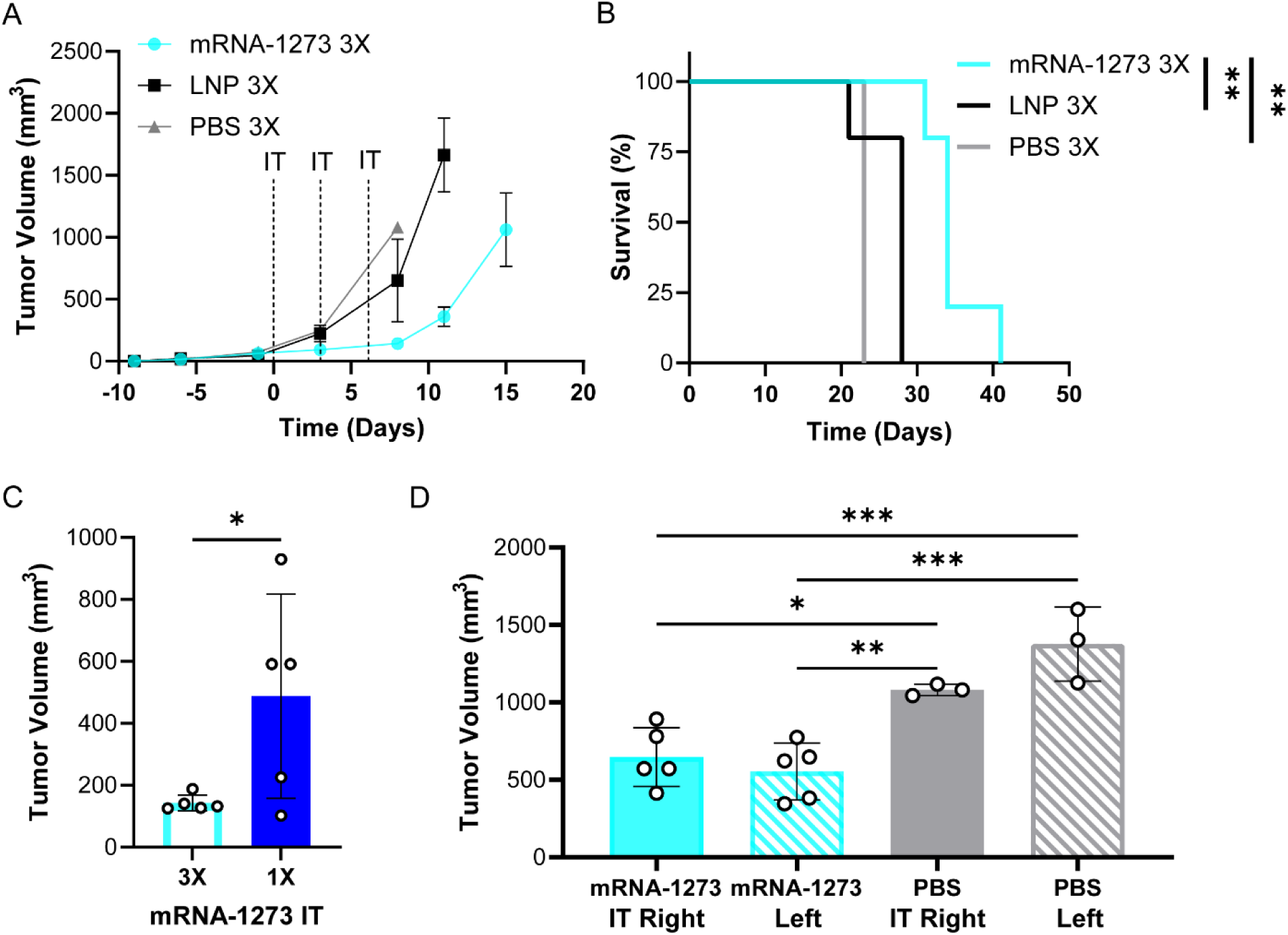
Enhanced tumor volume reduction results from using multiple mRNA-1273 IT injections. B16F10 (A) tumor growth curves and (B) survival curves following three 20 µL IT injections of 3 µg mRNA-1273, empty LNP or PBS (n = 5 mice treated with mRNA-1273 and LNP, n = 3 mice treated with PBS); hazard ration (HR) of mRNA-1273/LNP is 0.043 with log-rank (Mantel-Cox) test, **p = 0.0035, HR of mRNA-1273/PBS is 0.024 with log-rank (Mantel-Cox) test, **p = 0.0082 and HR of LNP/PBS is 0.16 with log-rank (Mantel-Cox) test, p>0.5. (C) B16F10 tumor volume measurements at 8 days post IT injection comparing three doses of mRNA-1273 versus a single mRNA-1273 IT dose (n = 5 mice/group). Abscopal effects of mRNA-1273 was evaluated using mice implanted with tumors on the right and left flanks. (D) B16F10 tumor volume measurements of treated right and untreated left flank tumors at day 8 post IT injection using three 20 µL IT injections of 3 µg mRNA-1273 or PBS (n = 5 mice/group). Data shown as mean ± SD; unpaired student t test or two-way ANOVA followed by Tukey’s multiple comparison test, \**p*<0.05, \*\**p*<0.01, \*\*\**p*<0.001.

### mRNA-1273 enhances immune checkpoint therapy response

We evaluated whether mRNA-1273 IT treatment synergized with immune checkpoint therapy (ICT). Four days post IT injection with mRNA-1273 or LNP, ICT treatment with anti-PD-1 and anti-CTLA-4 followed for three doses 3 days apart. A significant reduction in B16F10 tumor volume was observed from combination mRNA-1273 and ICT (Fig. 4A & S6A). It should be noted that combination PBS and ICT control group reached tumor endpoint, tumor diameter ≥ 15mm, before ICT treatments were completed (data not shown). A significant increase in survival was observed when B16F10 tumors were treated with combination mRNA-1273 and ICT (Fig. 4B). However, no difference in tumor growth curves or survival was observed between a single IT mRNA-1273 with or without ICT (Fig. S6B & C). We then combined multiple mRNA 1273 injections with ICT treatments starting 1 day post IT injection for a total of 3 treatments, and also observed a significant decrease in tumor volume from three doses of mRNA-1273 and ICT compared to combination LNP and ICT (Fig. 4C, S6D & E). A significant increase in survival time was observed from multiple treatments of combination mRNA-1273 and ICT (Fig. 4D). Multiple mRNA-1273 treatments further enhanced ICT efficacy as tumor growth was significantly reduced compared to multiple mRNA-1273 IT administration alone (Fig. S6F – H). In addition, the combination of multiple mRNA-1273 treatments with ICT increased survival time significantly compared to multiple mRNA-1273 treatments alone (Fig. S6I). Flow cytometry analysis of endpoint tumors from combination of multiple mRNA-1273 treatments with ICT did not significantly increase CD3^+^ cells or CD11b^+^ cells (Fig. S7A). A significant increase in CD8^+^ T cells was seen with approximately 25% of the CD8^+^ cells being IFNy^+^. Roughly 90% of the CD8^+^ T cells were also CD44^+^, with almost all of them being PD-1^+^. Interestingly, the endpoint tumors from combination of multiple LNP treatments with ICT had an increase in CD8^+^CD62L^+^, CD8^+^CD62L^+^PD-1^+^, CD8^+^CD44^+^CD62L^+^, and CD8^+^CD44^+^CD62L^+^ PD-1^+^ T cells when compared to mRNA-1273 group (Fig. S7B) There were no differences seen in the CD4^+^ T cells subsets (Fig. S7C). Interestingly, a single IT injection of mRNA-1273 followed by ICT treatment 1 day later did not demonstrate a significant delay in B16F10 tumor growth compared to PBS and ICT treatment (Fig. S6J & K). However, a single IT injection of mRNA-1273 followed by ICT treatment 1 day later did significantly delay YUMM1.G1 tumor growth compared to PBS with ICT (Fig. S6L). We additionally immunized mice with mRNA-1273 to first educate the immune system to the spike protein to see if we could additionally enhance melanoma reduction. Mice were immunized intramuscularly with mRNA-1273 at six weeks old. Two weeks post prime, B16F10 tumor inoculation occurred. Four weeks post prime (two weeks post B16F10 cell injection), intramuscular mRNA-1273 boost occurred to induce immune cell memory. One week following boost (three weeks post B16F10 cell injection), mRNA-1273 IT injection with and without ICT occurred. Immunization with mRNA-1273 did not enhance B16F10 tumor reduction using IT injection of mRNA-1273 or when combined with ICT (Fig. S8A). Immunization with or without ICT also did not increase long-term survival and has significantly increased tumor growth when compared to treatment without prime/boost (Fig. S8B – D).

**Figure 4.**
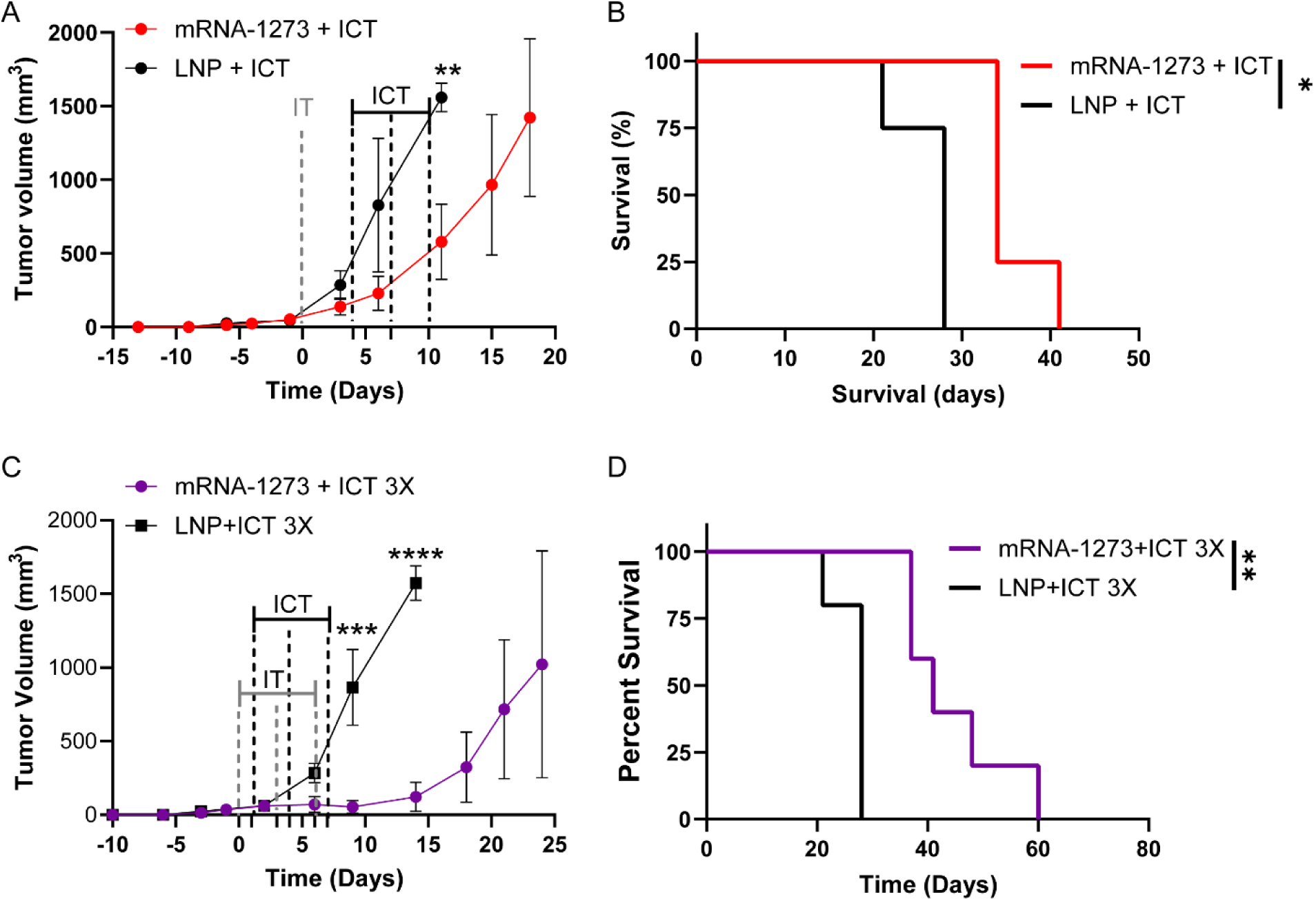
Synergistic effects of mRNA-1273 and immune checkpoint therapy (ICT). B16F10 (A) tumor growth curves and (B) survival curves following a single 20 µL IT injections of 3 µg mRNA-1273 or empty LNP combined with ICT 4 days later (n = 4 mice/group). PBS + ICT control group reached tumor endpoint before ICT treatments were completed. HR of mRNA-1273 + ICT/LNP + ICT is 0.050 with log-rank (Mantel-Cox) test, *p = 0.011. B16F10 (C) tumor growth curves and (D) survival curves following 3 doses of a 20 µL IT injections of 3 µg mRNA-1273 or empty LNP followed by ICT 1 day later (n = 5 mice/group); HR of mRNA-1273 + ICT 3X/LNP + ICT 3X is 0.043 with log-rank (Mantel-Cox) test, *p = 0.0039. Tumor growth data shown as mean ± SD, unpaired student t test comparing tumor volume post treatment, \*\**p*<0.01, \*\*\**p*<0.001, \*\*\*\**p*<0.0001.

### IT injection of mRNA-1273 does not result in systemic cytokine responses

Changes in cytokines in the tumor, lymph node, and sera at 48 h following an IT injection of mRNA-1273 was evaluated to determine if there were any systemic responses. Interestingly, no significant differences were observed between mRNA-1273 or PBS in the tumor, lymph node or sera among proinflammatory cytokines at the 48 h timepoint (Fig. 5A). Similarly, even after multiple doses of IT mRNA-1273, we did not measure seroconversion of anti-SARS-CoV-2 RBD IgG. Conversely, as expected we measured high titers of anti-RBD 2 weeks after a single dose of intramuscular administered mRNA-1273 (Fig. S9). This data suggests that IT injection of mRNA-1273 does not induce a systemic cytokine response.

**Figure 5.**
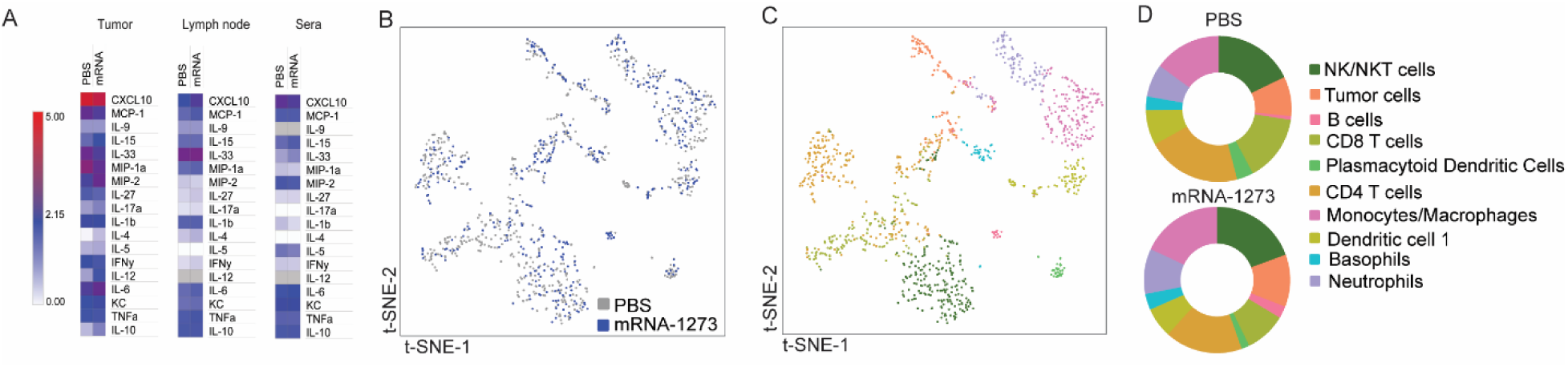
IT injection of mRNA-1273 does not alter cytokines or have transcriptional effects. (A) Cytokine levels of tumor, inguinal lymph node, and sera at 48 hours after IT injection with 3 µg mRNA-1273 or PBS. Heat map shown depicts mean pg/ml of each cytokine log10 transformed (n = 5). (B) tSNE plot of immune cell types separated by treatment groups. (C) tSNE plot of the cells defined by their single-cell transcriptome analysis from combined mRNA-1273 and PBS IT injections. (D) The proportions of cell projections from the tumors of mice with mRNA-1273 or PBS IT injections. (B-D) scRNAseq analysis was performed using pooled samples (n = 5), prior to library preparation.

### Evaluating transcriptional effects of mRNA-1273

The transcriptional effects of mRNA-1273 using single cell RNA sequencing (scRNA-seq) was also characterized. We observed the largest differences in tumor size at 4 days post IT injection, thus we hypothesized that cytotoxic CD8 T cell activity would be increased at this timepoint. Subcutaneous tumors from tumor-bearing mice either IT injected with mRNA-1273 or PBS were harvested, pooled, and live-cell sorted prior to library preparation and sequencing, to remove low-quality reads. As expected, mRNA-1273 tumor size and total cell numbers were reduced compared to tumors of PBS IT injected animals (Fig. S10). Our analysis found 1178 cells detected from the samples (Fig. 5B). We identified that 729 cells were from PBS IT injected mice and 449 were belonging to mRNA-1273 IT injected mice. IT injection of mRNA-1273 did not result in significant changes in cellular population clustering between PBS and mRNA-1273 treated samples at this timepoint (Fig. 5C-D). An analysis comparing the proportion of identified clusters did not reveal increases in CD8 T cells (Fig. 5D). Furthermore, we did not determine significant increases in proinflammatory genes such as *Cxcl10* (data not shown). Together the cytokine and transcriptomic analyses suggest that IT injection of mRNA-1273 does not induce a sustained proinflammatory response systemically or in the tumor microenvironment.

## Discussion

mRNA vaccines induced CD8^+^ T cell responses, prompting our investigation into how mRNA-1273 COVID-19 vaccine responses may impact tumor growth. Our study proposes that IT injection of mRNA-1273 reduces melanoma growth by enhancing CD8^+^ T cells in the tumor microenvironment, significantly prolonging survival. A single mRNA-1273 IT treatment significantly decreased tumor volume in two melanoma models, both of which are resistant to ICT. Specifically, our study shows that IT vaccination using mRNA-1273: 1) reduces tumor volumes on its own, 2) reduces tumors that are immune checkpoint therapy resistant, 3) improves anti-PD-1 and anti-CLTA-4 ICT outcomes.

Our findings reveal that B16F10 melanoma cells effectively internalize the mRNA-1273 vaccine and also exhibit robust expression of the encoded spike antigen. Expression of the spike antigen corresponded with high expression of C-X-C motif chemokine ligand 10 (CXCL10). Studies have demonstrated that T cell recruitment into tumor sites were predominately driven by CXCL10.^32–35^ Reduction in tumor volume occurred as early as 24 h post mRNA-1273 IT injection with a significant increase in CD8^+^ T cells into the tumor microenvironment. A major proportion of these intratumoral CD8^+^ T cells were activated CD8^+^GranzymeB^+^ T cells, which were significantly increased from mRNA-1273 treatment. While mRNA-1273 initially induced the generation of B16F10-specific CD8^+^TRP-2^+^ T cells at 24 h post IT injection, these melanoma-specific T subsets proved transient and were lost 4 days later. Despite initial suppression of tumor growth, long-term tumor control was not achieved following multiple mRNA-1273 treatments. Abscopal studies demonstrated that multiple mRNA-1273 IT treatments stimulated an immune response in untreated tumors within the same animal. However, once mRNA-1273 IT injections stopped, tumor outgrowth occurred. A caveat to these studies was that only a single B16F10 antigen, TRP-2, was evaluated to quantify a melanoma-specific response. This transient nature of melanoma-specific T cells following mRNA-1273 treatment combined with the minimal induced systemic immune changes may contribute to the observed lack of sustained, durable anti-cancer responses.

The remarkable clinical successes achieved through immunotherapy in improving the lives of cancer patients underscore the indispensable role of the immune system in cancer treatment. Yet, while immunotherapies have yielded notable successes, they have thus far provided benefits to only a fraction of patients. Tumors exhibiting substantial infiltration of CD8^+^ T cells have shown enhanced responsiveness to ICT.^8–13^ Our study evaluates the potential of mRNA-1273 as an adjuvant to enhance ICT outcomes. Enhanced responses from combination treatment with mRNA-1273 and ICT appears to be time dependent. Improved responses were only observed using multiple mRNA-1273 IT injections followed by ICT one day later for a total of 3 doses. No further tumor reduction was observed when ICT was started 4 days post mRNA-1273 in B16F10 tumors. The greater anti-tumor response of the multi-dose mRNA 1273 with ICT treatments started one day later, may be a result of the enhanced B16F10-specific CD8^+^ T cell infiltration that was observed at 24 h post mRNA-1273 IT injection. We recognize that our studies evaluated dual anti-PD1 and anti-CTLA-4 ICT treatment and given the known mechanistic differences in the activation of T cells between these immune checkpoint antibodies,^42^ future studies will evaluate responses separately.

Surprisingly, our data demonstrated a significant increase in CD3^+^ T cell subsets into the tumor and the draining lymph node following treatment with PBS compared to mRNA-1273 or LNP at 4 days post injection. It is possible that the injection of PBS into the tumor created a stress response to the injection itself. However, while we measured an increase of CD3^+^ T cells due to PBS injection compared to untreated tumors, we did not observe tumor suppression in the PBS injected mice. In addition to tumor suppression observed in mRNA-1273 treated mice, we measured a significant increase in TRP-2 specific cells only in tumors of mRNA-1273 injected samples. Therefore, while the stress response due to PBS injection may have induced a measurable proinflammatory cytokine response, this did not translate to tumor suppression as we observed with mRNA-1273 treatment.

scRNAseq data did not demonstrate a significant difference in the cellular populations recruited to the tumors. However, given that tumors were significantly smaller 4 days post IT injection with mRNA-1273, immune cell driven changes to the tumor microenvironment may have occurred at earlier timepoints and may be one explanation for the lack of transcriptional differences observed. For example, we observed in our intravital imaging studies that at 48 h post IT mRNA-1273 treatment, areas where the tumor had been eradicated, as evidenced by minimal GFP fluorescence, had minimal detectable CD8^+^ T cells which were detected 24 h earlier. Furthermore, the scRNAseq analysis removed cells with low quality read data, and before RNA sequencing we removed any cells positive for 7-AAD viability stain. It is possible that to do these quality constraints some of the changes in the tumor transcriptome could have been overlooked. Future studies with mRNA vaccines should evaluate early changes such as 2 – 6 h since LNPs have been shown to be internalized and express antigens by cells within those time frames.

Given the lack of long-term durable responses, re-challenges studies to demonstrate the presence of memory T cells could not be conducted. Surprisingly, seroconversion of the mRNA-1273 antigen was not observed, as evident by anti-RBD ELISA following IT administration. Additionally, the transient presence of B1610-specific T cells limited our studies to subcutaneous tumor models. However, our studies demonstrate a significant reduction in tumor volume from IT mRNA-1273 injection with corresponding enhanced infiltration of activated CD8^+^ T cells into the tumor microenvironment. These data suggest that expression of spike protein by tumor cells were not sufficient to elicit durable B16F10-specific T cell responses. In our present study, we are implementing an exogenous antigen, and we recognize that our studies are limited to the absence of a tumor-associated antigen. The exclusion of a B16F10 specific antigen with our mRNA-1273 treatments may fail to generate tumor-specific T cells, which are required to induce a memory response. Studies are underway to evaluate combination mRNA-1273 with B16F10-specific vaccines to enhance both the infiltration and activation of melanoma-specific T cells into the tumor microenvironment. It is possible that by polarizing the immune response during priming with a therapeutic tumor-specific vaccine, further recruitment of CD8^+^ T cells to the tumor will occur. These findings have important implications for mRNA vaccines in priming patients to respond to existing immunotherapies.

## Materials and Methods

### Cells

B16F10 and YUMM1.G1 melanoma cells were purchased from the American Type Culture Collection (ATCC, VA, USA). B16F10 click beetle green luciferase and green fluorescent protein (CBG-GFP) cells were previously described.^26, 37, 39, 40^ B16F10 cells were cultured in DMEM supplemented with 10% heat-inactivated fetal bovine serum (FBS). YUMM1.G1 were cultured in DMEM:F12 supplemented with 10% heat-inactivated FBS and 1% NEAA. Cell cultures were grown at 37°C in a humidified 5% CO_2_ atmosphere. All cell lines tested negative for mycoplasma.

### In vitro studies

2 × 10^5^ B16F10 cells were seeded in a 6-well plate. B16F10 cells were cultured up to 96 hours after inoculation with 40 μg of mRNA-1273 SARS-CoV-2 vaccine. Cell culture supernatants and cell lysates were collected every 24 hours. Lysates were prepared with RIPA Buffer (Thermo Fisher) and HALT proteinase inhibitor (Thermo Fisher Scientific). SARS-CoV-2 Spike protein production was confirmed by western blot of lysates. Cell lysate concentrations were normalized by total protein concentration. Western blots were prepared using Bolt 4-12% Bis-Tris Plus Protein Gels (Invitrogen) and transferred to PVDF membrane using iBlot 2 transfer. Western blot membrane was blocked using 3% BSA in tris-phosphate buffer. Spike protein was confirmed by sera from convalescent SARS-Cov-2 challenged mice, and Spike-S1 subunit (Sino Biological part no. 40591-V08H).

### *In vivo* subcutaneous tumor model

The Institutional Animal Care and Use Committee at West Virginia University approved all animal protocols (2109047227). 5 × 10^4^ B16F10 cells were injected subcutaneously on the right flank of female 8-week-old C57BL/6 (Charles River Laboratories, NY, USA)) mice. Before tumor cell injection, the fur was removed on the right flank. Tumors were measured by calipers every 3-4 days once palpable. Following institutional animal guidelines, animals were euthanized once tumors reached 1.5 cm in diameter or were ulcerated greater than 0.5 cm. When tumors reached 5 mm in diameter, an IT injection of 20 μL of 3 μg mRNA-1273, empty LNP or PBS control occurred. mRNA-1273 (Spikevax, Moderna) vaccine was obtained from West Virginia United Hospital. Empty LNP without mRNA was prepared using the using the flash nanoprecipitation approach as described elsewhere.^43^ The composition of empty LNP is similar to that of mRNA-1273 lipid nanoparticles. Empty LNP were 120 nm in size when analyzed using dynamic light scattering. Animals were treated with ICT (1^st^ dose was 200 µg Anti-CTLA-4 and 200 µg anti-PD-1 followed by 2 doses of 100 µg Anti-CTLA-4 and 100 µg anti-PD-1 administered intraperitoneally every 3 days). Anti-CTLA-4 (9D9), and anti-αPD-1 (RMP1-14) were obtained from Bio X cell (Bio X cell, NH, USA).

### Flow Cytometry

At 24 h following a 50 μL injection of 50 μg mRNA-1273 vaccine or PBS control using an intramuscular injection, 8-week old C57BL/6 (Charles River Laboratories, NY, USA)) mice were euthanized by pentobarbital injection. The muscle where the site of injection occurred and the inguinal lymph node were analyzed by flow cytometry. At various timepoints, tumor-bearing mice were euthanized by carbon dioxide asphyxiation. Single cell suspension of cells were harvested from the spleen, lymph nodes, and tumor. Tumors and muscle were dissociated using a mouse tumor dissociation kit, or skeletal muscle dissociation kit, respectfully (Miltenyi Biotec, CA, USA). Red blood cells were lysed using 1X red blood cell lysis buffer (Thermo Fisher Scientific, CA, USA). Remaining cells were resuspended in cell staining buffer (BioLegend, CA, USA ) at a concentration of 2 × 10^5^ – 1 × 10^6^ cells per 100 µl. Fc receptors are blocked using 10 µg/ml ChromPure of mouse IgG (Jackson ImmunoResearch Inc., PA, USA) per 10^6^ cells in a 100 µl volume for 10 minutes on ice. Cells were washed with cell staining buffer and incubated with antibody mix (Table S1) for 15 – 20 minutes on ice in the dark. Cells were fixed using 200 µl Fixation buffer (Thermo Fisher Scientific, CA, USA) for 20 min at 4°C in the dark. Cells are washed 2 times with 1X PermWash (BD Bioscience, CA, USA) and permeabilized using Permeabilization solution (BD Bioscience, CA, USA). Intracellular antibodies (Table S1) were incubated for 20 minutes at room temperature in the dark. Stained cells were analyzed using the Cytek® Aurora (Cytek, MD, USA) within 2 weeks of staining. Data was analyzed using FCS express (De Novo Software, CA, USA).

### Intravital Imaging

Skin window chamber implantation, imaging and analysis were previously described using 8-week old male and female C57BL/6 mice.^40 41^ Briefly, following skin window chamber implantation and B16F10 cell inoculation, microscopy occurred on day 2 post implantation. Skin window chamber microscopy was done using a Nikon confocal AX microscopy (Nikon, NY, USA). 50 μL of 0.15 µg CD8a-APC (Miltenyi, CA, USA) antibody in PBS was administered through retro-orbital injection. Microscopic imaging occurred 30 min post antibody injection using 3 fields of view per mouse at 20X objective. B16F10 tumors were visualized within the skin window chamber using GFP. Following background imaging, an IT injection of 10 μL of 3 μg mRNA-1273 or PBS control occurred. 24h post injection, intravital microscopy occurred again. Images were analyzed using the Nikon Elements software (Nikon, NY, USA) using the automated measurement analysis macro. An individual threshold was applied to each imaging channel and positive CD8a-APC positive cells were quantified using the following automated measurement parameters; size: 5 µm to 10 µm, circularity: 0 to 1, smooth: 5X, fill holes: on and exclude objects touching frame borders. The average of the 3 fields of view from each imaging time point for each animal was calculated.

### Cytokine analysis

Blood draws by submandibular bleed were performed on subcutaneous tumor-bearing mice at various timepoints during treatment and stored at -80° C until analysis. Serum was prepared using serum separator microtiter tubes (BD). Cytokines were analyzed using a V-Plex Mouse Cytokine 19-Plex kit (Mesoscale Diagnostics), following the manufacturer’s protocol.

### Single cell RNA sequencing

At 4 days post intratumor injection of mRNA-1273 or PBS, mice were euthanized by carbon dioxide asphyxiation. Tumors were smashed and washed with PBS through a 70um cell strainer to achieve a single-cell suspension. Samples were pooled per group and combined at equal proportion per individual sample. Single-cell suspensions were stained with 7-AAD (STEMCELL Technologies), and viable cells sorted using BD Aria. RNA was isolated using Chromium Next GEM Single Cell 5’ Kit (10X Genomics) following manufacturers protocol. Read coverage was calculated at 2 billion paired reads per 5’ library samples. scRNAseq analysis was conducted using Loupe Browser (7.0.1). Low quality reads were filtered using the following thresholds: Unique molecular identifiers (UMI) 500-50,000; genes per barcode 300-7000; and less than 20% mitochondrial UMIs. The cells lacking Ptprc gene expression assigned as tumor cells. To assign major immune cell types to each cluster, we annotated each cluster based on its top 10 upregulated, differentially expressed genes compared to other clusters. Top 10 upregulated genes in each cell cluster were shown in Supplementary Figure S11.

### Quantification of anti-SARS-CoV-2 RBD IgG antibodies

Serum was collected after the final timepoint (∼20 days from initial dose) from the multiple (3X) mRNA-1273 IT (3 µg) treated mice. Non-tumor bearing six-week-old, C57B6 mice received one IM dose of 5 µg mRNA-1273. After 14 days serum was collected from submandibular bleeds. Anti-SARS-CoV-2 RBD IgG levels were quantified using ELISA^44^ from sera of the two cohorts. High-binding microtiter plates were coated with 2 µg/ml of WA-1 S RBD (Sino Biological), and incubated for 24 h at 4 °C. The next day plates were washed 3 times with wash buffer (0.1 % Tween-20+PBS). Plates were blocked using 3% nonfat milk in wash buffer and incubated at room temperature for 1 h. After 3X washes, sera were serial diluted in 1% nonfat milk and wash buffer, using 2-fold dilutions starting at 1:50 dilution. Samples were incubated for 1 h at room temperature while shaking. Plates were then washed 4X, and secondary antibody added (goat anti-mouse IgG HRP, Novus Biological NB7539; 1:2000) and incubated for 1 h while shaking at room temperature. Following 5X wash, plates were developed using TMB substrate (Biolegend) for 15 mins. Reaction was stopped using 2N sulfuric acid and absorbance read at 450nm on a Synergy HI reader. Serum antibody levels were quantified using area under curve analysis in GraphPad prism.

### Statistical analyses

Graphs were made and statistical analyses were performed using GraphPad Prism V10.1.0 (GraphPad Software, Inc, CA, USA). Data were expressed as mean ± SD. For analysis of three or more groups, analysis of variance (ANOVA) tests were performed followed by a Tukey’s or Bonferroni’s multiple comparisons test, or false discovery rate test. Standard comparison of survival curves used the log rank (Mantel-Cox) analysis. Analysis of differences between two normally distributed paired test groups were performed using a Student’s t-test. P values were considered statistically significant if *p* < 0.05.

## Author contributions

DB and KML performed all *in vitro* and *in vivo* experiments, quantified and analyzed the data, provided critical intellectual input, and wrote and edited the manuscript. ESK performed the bioinformatics analysis of the scRNAseq data and edited the manuscript. KSL performed immunization studies and edited the manuscript. SB and BM provided empty LNP for control studies. FHD supervised the study, quantified and analyzed the data, provided critical intellectual input, and edited the manuscript. TWL supervised the study, quantified and analyzed the data, provided critical intellectual input, wrote and edited the manuscript.

## Data Availability Statement

All data relevant to the study are included in the article, uploaded as supplementary information or will be deposited in the public, open access repository NCBI SRA/GEO

