## Supplemental File for "Intratumoral administration of mRNA-1273 vaccine delays melanoma growth in mice"

**Running Title:** mRNA-1273 vaccine delays melanoma

**Authors:** Dylan T. Boehm[†,1,2], Kaitlyn M. Landreth[†,1], Emel Sen Kilic[1,2], Katherine S. Lee[1,2],

Bishal Misra[3], Sharan Bobbala[3], F. Heath Damron[1,2], Tracy W. Liu[1,4] *

**Affiliations:**

[1] Department of Microbiology, Immunology, and Cell Biology, West Virginia University, Morgantown, WV, 26506, USA.

[2] Vaccine Development Center at West Virginia University Health Sciences Center, Morgantown, WV, United States

[3] Department of Pharmaceutical Sciences, West Virginia University, Morgantown, WV, 26506, USA.

[4] WVU Cancer Institute, West Virginia University, Morgantown, WV, 26506, USA.

[†]Authors contributed equally

*Corresponding author:

Tracy W. Liu, Ph.D.,
Department of Microbiology, Immunology, and Cell Biology
School of Medicine
West Virginia University
64 Medical Center Drive
Morgantown, WV, 26506

ORCID ID: 0000-0003-0671-8390

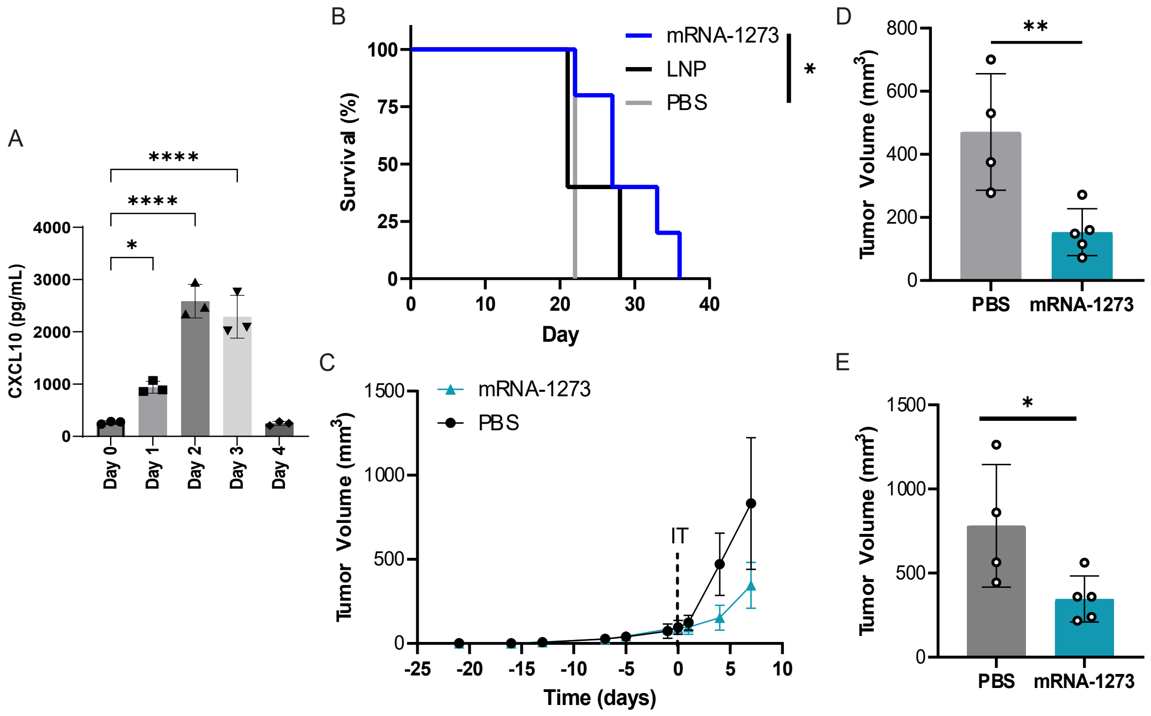


Figure S1. Intratumor injection of mRNA-1273 delays melanoma growth. (A) CXCL10 concentration in supernatant of B16F10 cells treated with 40 ug mRNA-1273 from day 0 – 4. (B) Survival curve of B1610 subcutaneous tumors administered a 20 μL intratumoral injection (IT) of 3 μg mRNA-1273, LNP or PBS (n = 5 mice/group); log-rank (Mantel-Cox) test *p<0.05. (C) YUMM1.G1 tumor growth curves and tumor volume measurements at (D) 4 days and (E) 7 days following a 20 μL IT of 3 μg mRNA-1273 or PBS (n = 4 – 5 mice/group). (A) Data shown as mean ± SD; ANOVA followed by Dunnett's multiple comparison test, *p<0.05, ****p<0.0001. (B-E) Day 0 represents time of IT injection when tumors reached 5 mm in diameter. Data shown as mean ± SD; Student t test, *p<0.05.

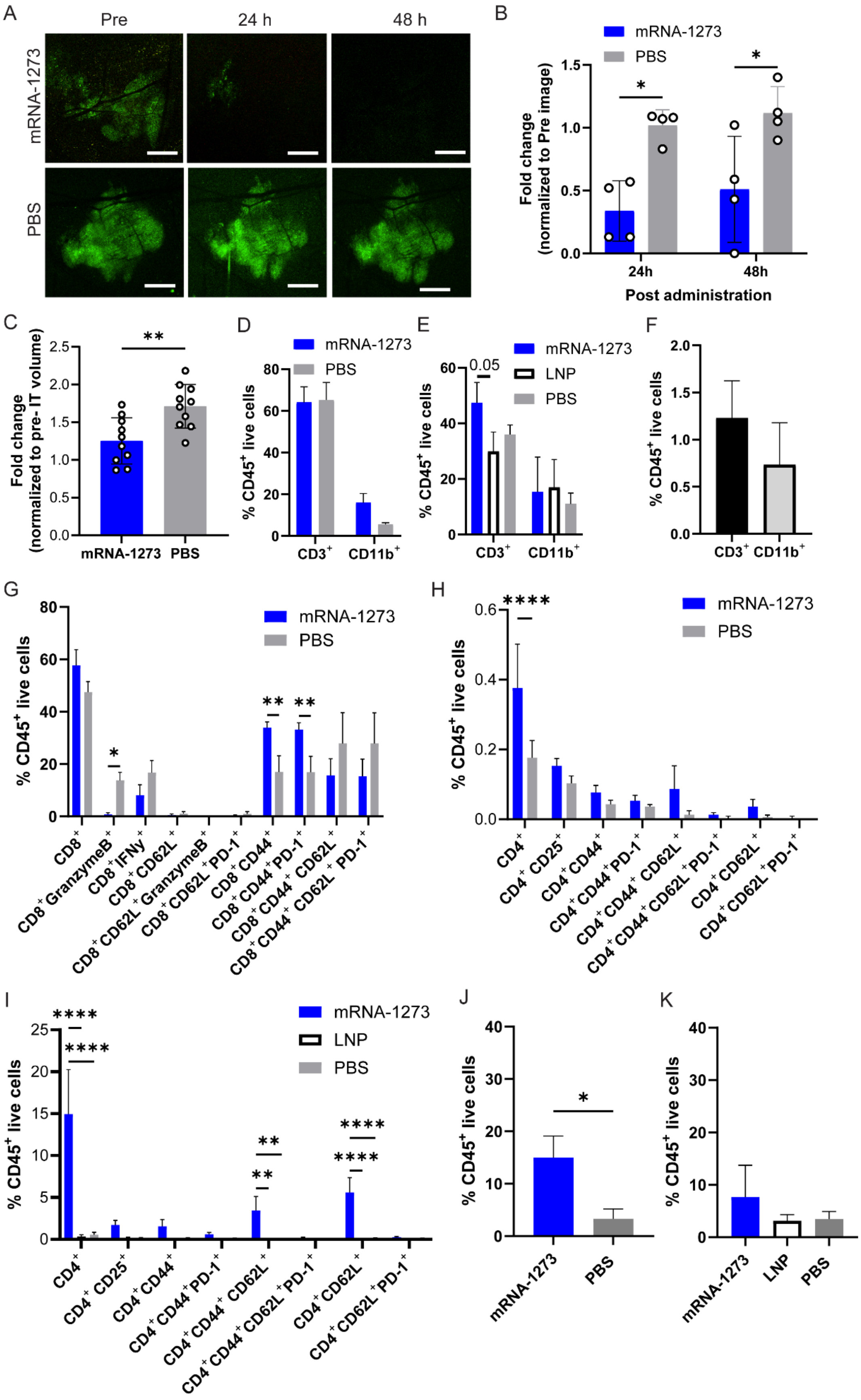
A
Pre
24 h
48 h
mRNA-1273
PBS
B
mRNA-1273
PBS
Fold change (normalized to Pre image)
24h
48h
Post administration
C
Fold change (normalized to pre- IT volume)
mRNA-1273
PBS
D
mRNA-1273
PBS
% CD45+ live cells
CD3+
CD11b+
E
0.05
mRNA-1273
LNP
PBS
% CD45+ live cells
CD3+
CD11b+
F
% CD45+ live cells
CD3+
CD11b+
G
mRNA-1273
PBS
% CD45+ live cells
CD8+
CD8+ GranzymeB+
CD8+ IFNγ+
CD8+ CD62L+
CD8+ CD62L+ GranzymeB+
CD8+ CD62L+ PD-1+
CD8+ CD44+
CD8+ CD44+ PD-1+
CD8+ CD44+ CD62L+
CD8+ CD44+ CD62L+ PD-1+
H
mRNA-1273
PBS
% CD45+ live cells
CD4+
CD4+ CD25+
CD4+ CD44+
CD4+ CD44+ PD-1+
CD4+ CD44+ CD62L+
CD4+ CD44+ CD62L+ PD-1+
CD4+ CD62L+
CD4+ CD62L+ PD-1+
I
mRNA-1273
LNP
PBS
% CD45+ live cells
CD4+
CD4+ CD25+
CD4+ CD44+
CD4+ CD44+ PD-1+
CD4+ CD44+ CD62L+
CD4+ CD44+ CD62L+ PD-1+
CD4+ CD62L+
CD4+ CD62L+ PD-1+
J
% CD45+ live cells
mRNA-1273
PBS
K
% CD45+ live cells
mRNA-1273
LNP
PBS

Figure S2. mRNA-1273 vaccine intratumoral treatment reduces tumor growth and alters tumor-immune composition. (A) Representative intravital image of tumor (GFP) prior to (Pre), 24 h and 48 h post 10 µL IT injection of 1.5 µg mRNA-1273 or PBS; 2X objective, scale bar represents 1 mm. (B) Corresponding fold change in tumor size quantified using tumor GFP area normalized to Pre-image; a fold change < 1 means tumor size has decreased (n = 4 mice/group). (C) Fold change in tumor volume at 24 h post IT injection of 20 µL IT injection of 3 µg mRNA-1273 or PBS (n = 10 mice/group). Flow cytometry analysis of $CD3^+$ and $CD11b^+$ subsets at (D) 24 h and (E) 4 days post IT injection of mRNA-1273, PBS or LNP and (F) at endpoint in untreated B16F10 tumors (n = 3 mice/group) . Flow cytometry analysis of (G) $CD8^+$ and (H) $CD4^+$ T cell subsets within the tumor at 24 h, (I) $CD4^+$ T cell subsets within the tumor at 4 days post IT injection and $CD8^+TRP2^+$ T cells at (J) 24 h and (K) 4 days post IT injection (n = 3 mice/group). Data shown as mean ± SD; one-way or two-way ANOVA followed by Tukey's multiple comparison test, *p<0.05, **p<0.01, ***p<0.001, ****p<0.0001.

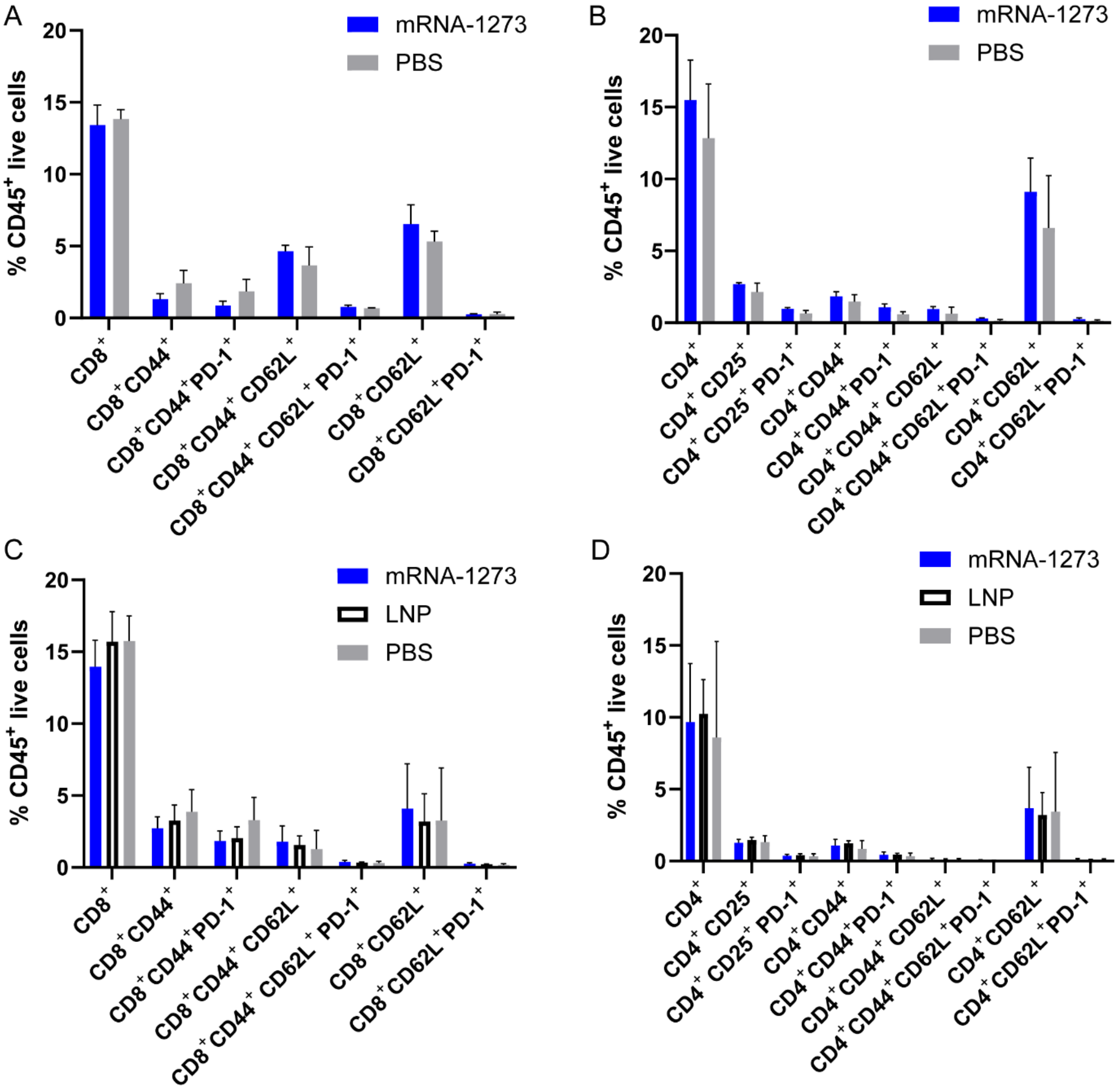


Figure S3. No differences in T cell subsets were observed in the spleen following IT injection. Flow cytometry analysis of the spleen showing (A) $CD8^+$ and (B) $CD4^+$ T cell subsets at 24 h post IT injection and (C) $CD8^+$ and (D) $CD4^+$ T cell subsets at 4 days post IT injection (n = 3 mice/group). Data shown as mean ± SD; one-way or two-way ANOVA followed by Tukey's multiple comparison test, $p>0.05$ for all comparisons.

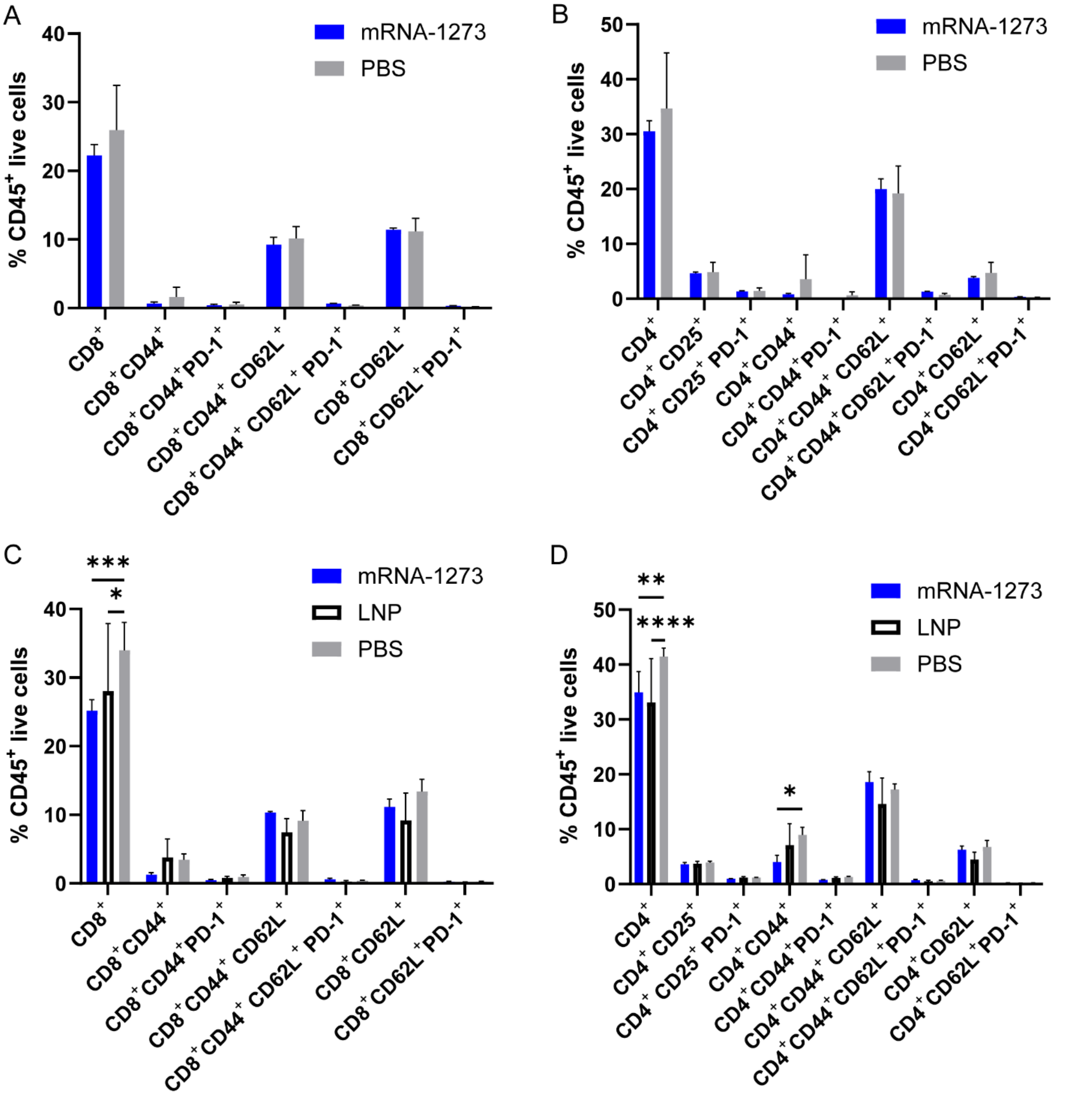


Figure S4. T cell subset analysis of draining lymph node following IT injection. Flow cytometry analysis of the lymph node showing (A) $CD8^+$ and (B) $CD4^+$ T cell subsets at 24 h post IT injection and (C) $CD8^+$ and (D) $CD4^+$ T cell subsets at 4 days post IT injection (n = 3 mice/group). Data shown as mean ± SD; one-way or two-way ANOVA followed by Tukey's multiple comparison test, *$p<0.05$, **$p<0.01$, ***$p<0.001$, ****$p<0.0001$.

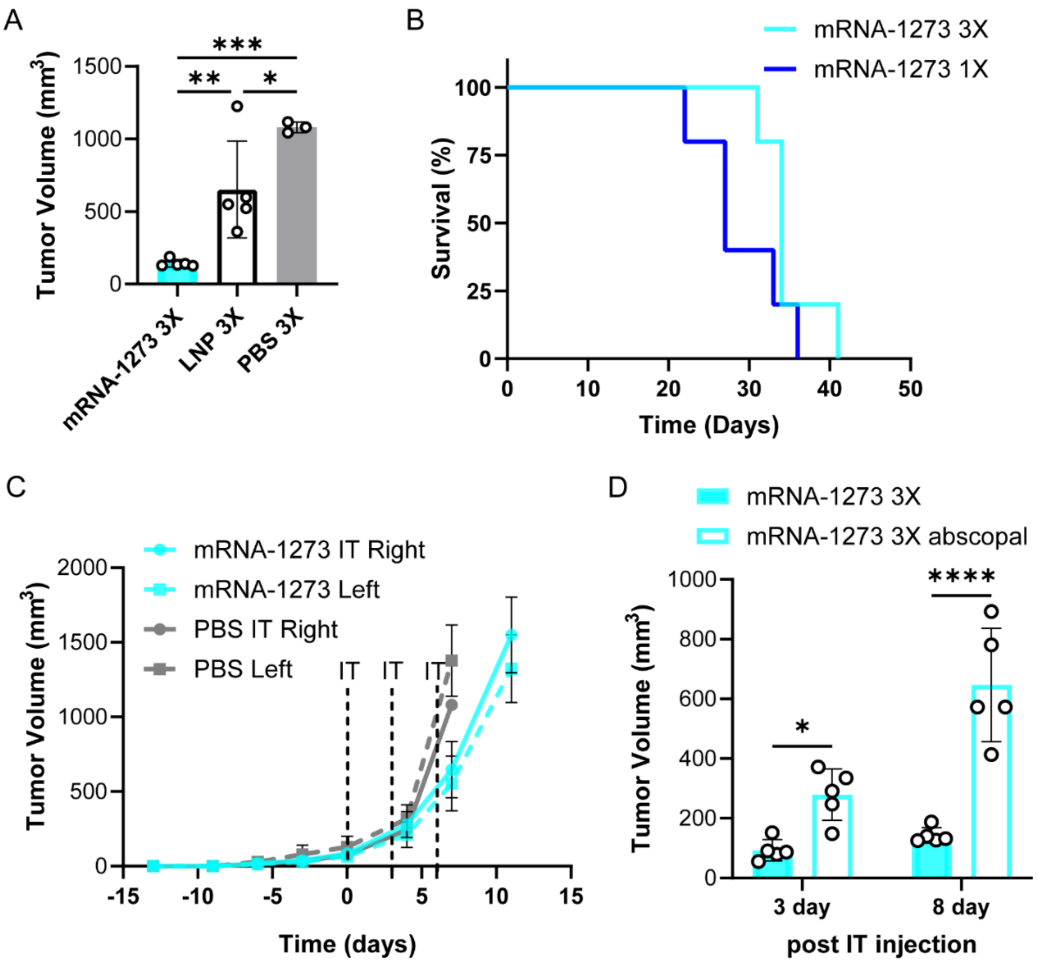


Figure S5. Multiple IT treatments of mRNA-1273 reduces B16F10 tumor volume. (A) B16F10 tumor volume at 8 days post injection following three 20 µL IT injection of 3 µg mRNA-1273, LNP or PBS (n = 5 mice treated with mRNA-1273 and LNP, n = 3 mice treated with PBS). (B) Survival curve comparing B16F10 tumor-bearing mice treated with a single dose (1X) or three doses (3X) of 20 µL IT injection of 3 µg mRNA-1273 (n = 5 mice/group); log-rank (Mantel-Cox) test $p>0.05$. (C) Tumor volume measurements following three 20 µL IT injections of 3 µg mRNA-1273 or PBS of treated right and untreated left flank tumors (n = 5 mice/group). (D) B16F10 treated tumor volume measurements comparing three doses of mRNA-1273 in mouse bearing a single tumor or two tumors on the right and left flank (abscopal) at day 3 and 8 post IT injection Data shown as mean ± SD; one-way or two-way ANOVA followed by Tukey's multiple comparison test, *$p<0.05$, **$p<0.01$, ***$p<0.001$, ****$p<0.0001$.

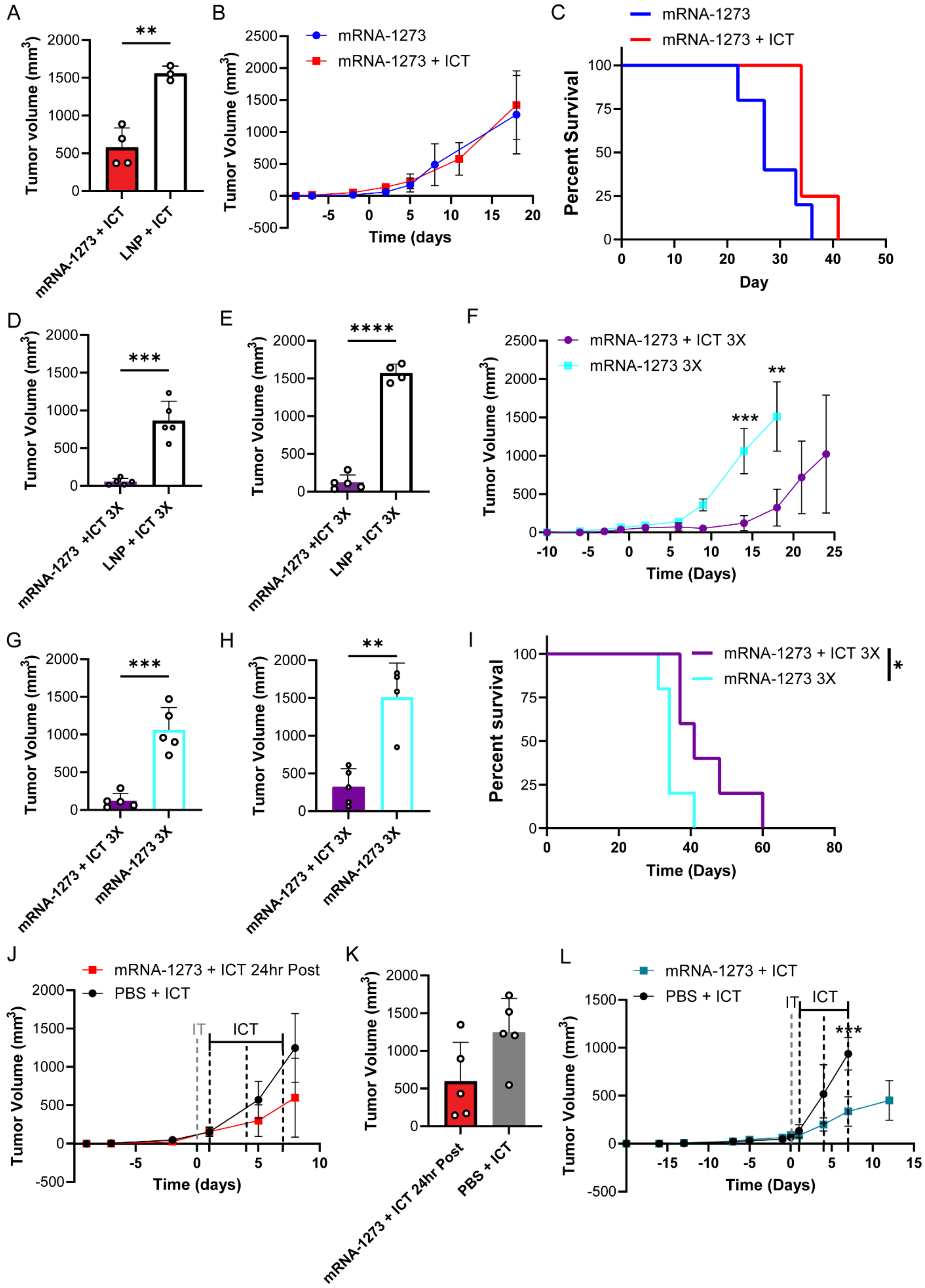

A
Tumor volume (mm3)
**
mRNA-1273 + ICT
LNP + ICT
B
mRNA-1273
mRNA-1273 + ICT
Tumor Volume (mm3)
Time (days
C
mRNA-1273
mRNA-1273 + ICT
Percent Survival
Day
D
Tumor Volume (mm3)
***
mRNA-1273 +ICT 3X
LNP + ICT 3X
E
Tumor Volume (mm3)
****
mRNA-1273 +ICT 3X
LNP + ICT 3X
F
mRNA-1273 + ICT 3X
mRNA-1273 3X
Tumor Volume (mm3)
***
**
Time (Days)
G
Tumor Volume (mm3)
***
mRNA-1273 + ICT 3X
mRNA-1273 3X
H
Tumor Volume (mm3)
**
mRNA-1273 + ICT 3X
mRNA-1273 3X
I
mRNA-1273 + ICT 3X
mRNA-1273 3X
*
Percent survival
Time (Days)
J
mRNA-1273 + ICT 24hr Post
PBS + ICT
Tumor Volume (mm3)
IT
ICT
Time (days)
K
Tumor Volume (mm3)
mRNA-1273 + ICT 24hr Post
PBS + ICT
L
mRNA-1273 + ICT
PBS + ICT
IT
ICT
***
Tumor Volume (mm3)
Time (Days)

Figure S6. Combination mRNA-1273 and ICT treatments. (A) B16F10 tumor volume at 11 days post injection following a single 20 µL IT injection of 3 µg mRNA-1273 or LNP followed with ICT 4 days later (n = 4 mice/group). B16F10 (B) tumor growth curve and (C) survival curve comparing a single IT injection of mRNA-1273 without and with ICT 4 days later (n = 4/5 mice/group). B16F10 tumor volume at (D) 9 and (E) 14 days post three doses of a 20 µL IT injection of 3 µg mRNA-1273 or LNP followed with ICT a day later (n = 5 mice/group). B16F10 (F) tumor growth curves, tumor volume at (G) 14 and (H) 18 days post three 20 µL IT injection of 3 µg mRNA-1273 with or without ICT and (I) survival curves (n = 5 mice/group). B16F10 (J) tumor growth curve and (K) tumor volume at day 7 post a single 20 µL IT injection of 3 µg mRNA-1273 or PBS followed with ICT 1 day later (n = 5 mice/group). (L) YUMM1. tumor growth curve following treatment with a single 20 µL IT injection of 3 µg mRNA-1273 or PBS followed with ICT 1 day later (n = 5 mice/group). Data shown as mean ± SD; unpaired student t test, *p<0.05, **p<0.01, ***p<0.001, ****p<0.0001.

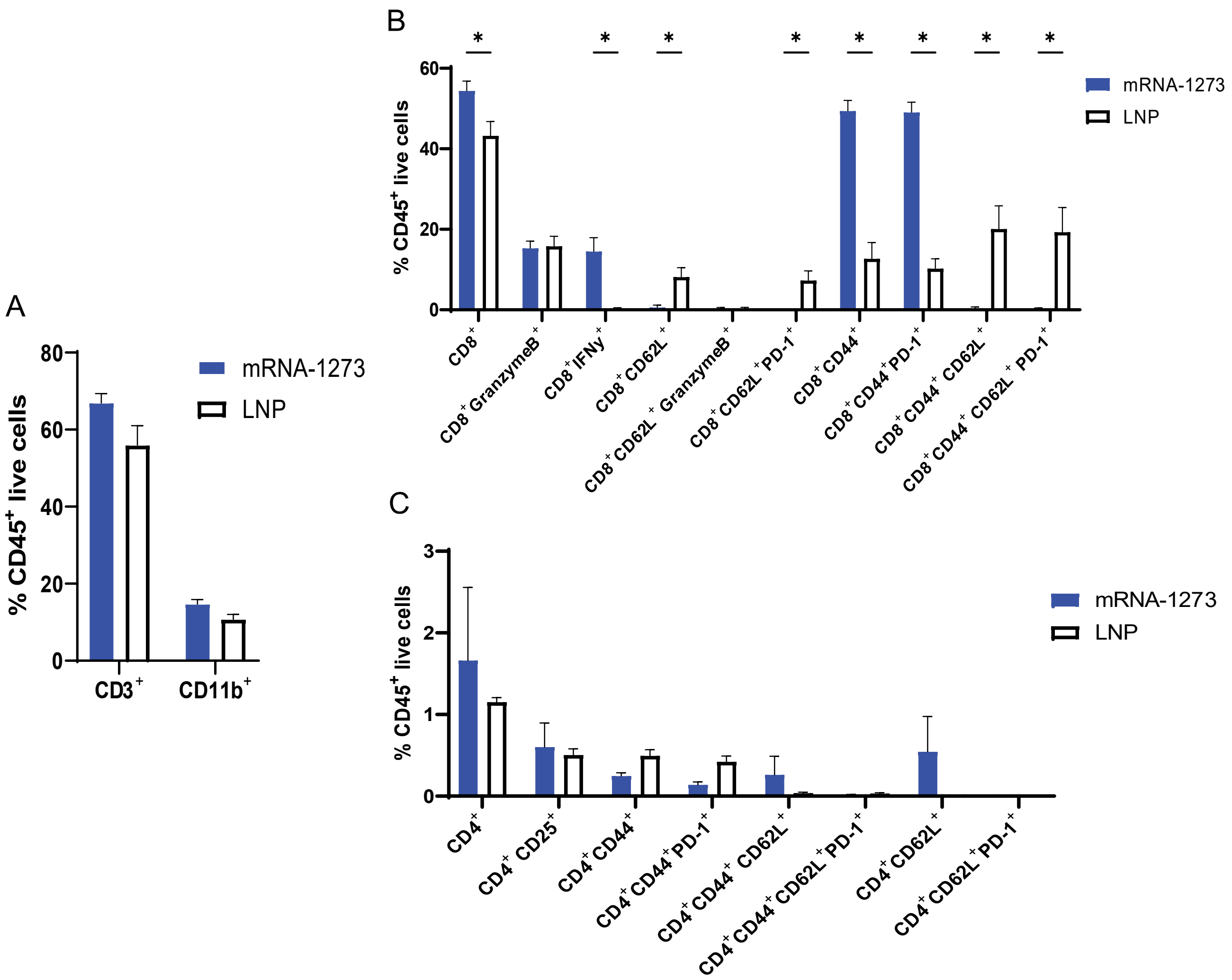


Figure S7. mRNA-1273 vaccine intratumoral treatment alters tumor-immune composition at endpoint of tumors. (A) Flow cytometry analysis of $CD3^+$ and $CD11b^+$ subsets post injection of mRNA-1273, PBS or LNP (n=3 mice per group). (B) Flow cytometry analysis of $CD8^+$ and (H) $CD4^+$ T cell subsets within the tumor at endpoint (n = 3 mice/group). Data shown as mean ± SD; one-way or two-way ANOVA followed by Tukey's multiple comparison test, *p<0.05, **p<0.01, ***p<0.001, ****p<0.0001.

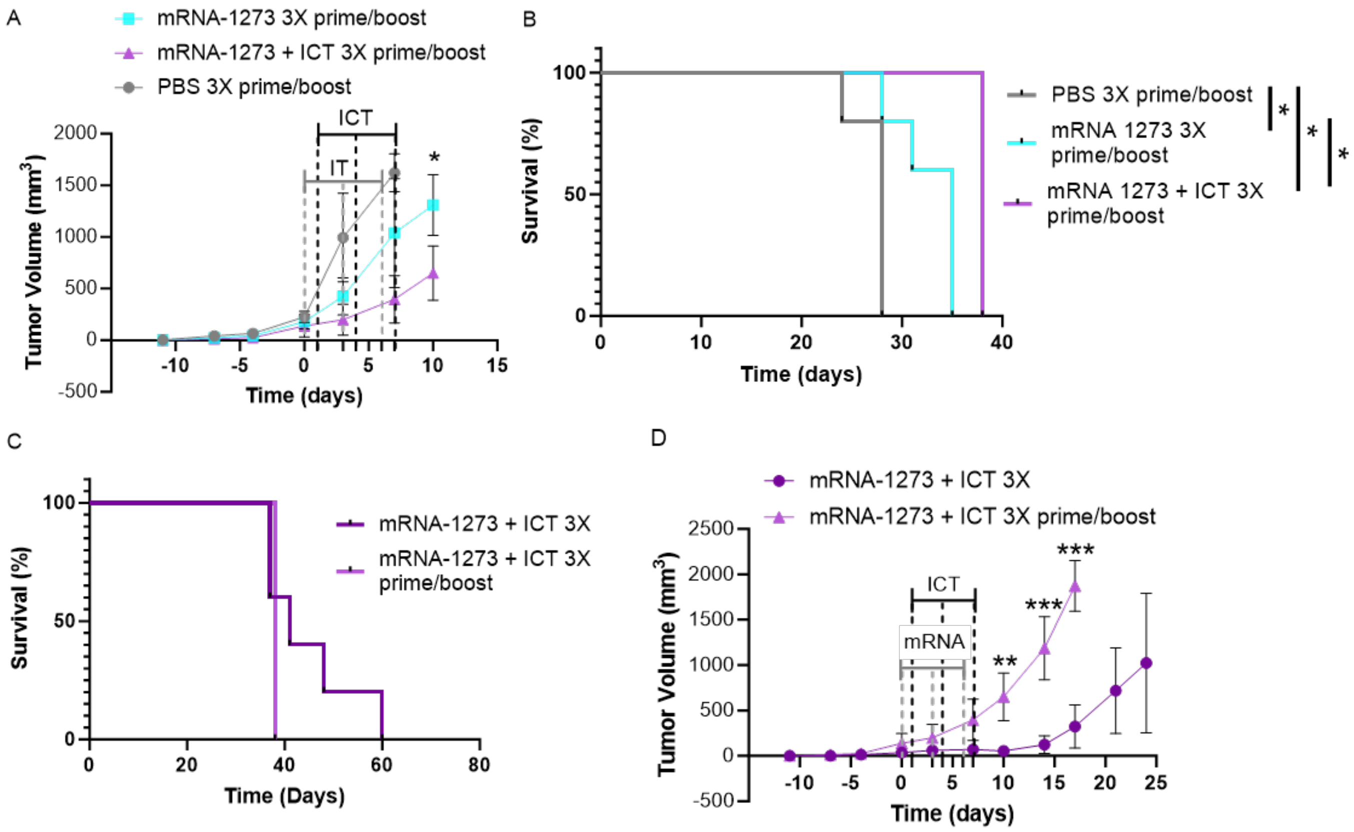


Figure S8. Intramuscular prime and boost with mRNA-1273 prior to intratumoral injection does not enhance anti-tumor response. (A) B16F10 tumor growth curves following a single 20 µL IM injection of 3 µg mRNA-1273 or PBS followed with a 20 uL IM boost 4 weeks post prime and a 20 uL IT injection when tumors reached 5mm in diameter (n = 3/5 mice/group). (B) survival curve comparing prime/boost PBS, mRNA-1273 and mRNA1273+ICT, log-rank (Mantel-Cox) test *p<0.05 (n = 3/5 mice/group). (C) survival curve comparing mRNA-1273 + ICT 3X to prime/boost mRNA1273+ICT 3X, log-rank (Mantel-Cox) test *p<0.05 (n = 3/5 mice/group). (D) B16F10 tumor growth curves comparing mRNA-1273+ICT 3X and mRNA-1273+ICT 3X prime/boost. Data shown as mean ± SD; unpaired student t test, *p<0.05, **p<0.01, ***p<0.001, ****p<0.0001.

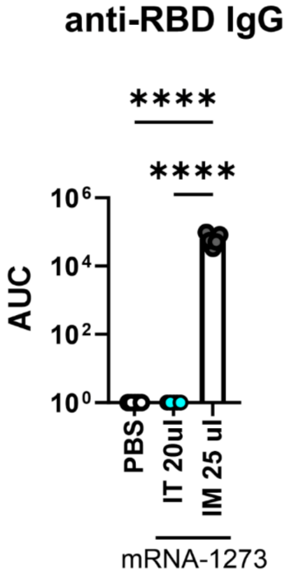


Figure S9. Anti- SARS-CoV-2 RBD IgG antibody levels after IM and IT mRNA-1273 injections. Anti-RBD antibody levels in serum of mice treated with 3X - mRNA-1273 IT (3 µg) at 20 days post first injection compared to mice treated with single mRNA-1273 IM (5µg) 14 days post injection. Data shown as mean ± SD; ANOVA followed by Tukey's multiple comparison test, ****p<0.0001.

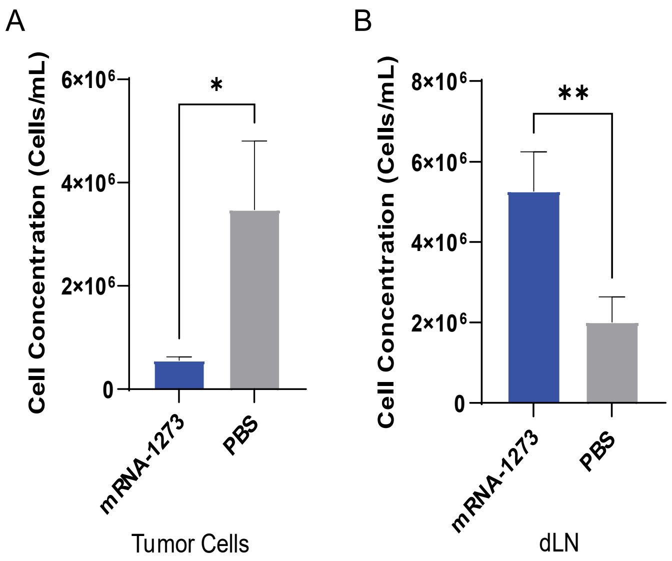


Figure S10. Single dose of mRNA-1273 reduces tumor cell number and increases cell number in draining lymph node (dLN) at single-cell time point. (A) Number of cells isolated from tumors 4 days post IT injection of 20 uL or 3ug of mRNA-1273 or PBS. (B) Number of cells isolated from dLN 4 days post IT injection of 20 uL or 3ug of mRNA-1273 or PBS. Data shown as mean ± SD; Student t test, *p<0.05.

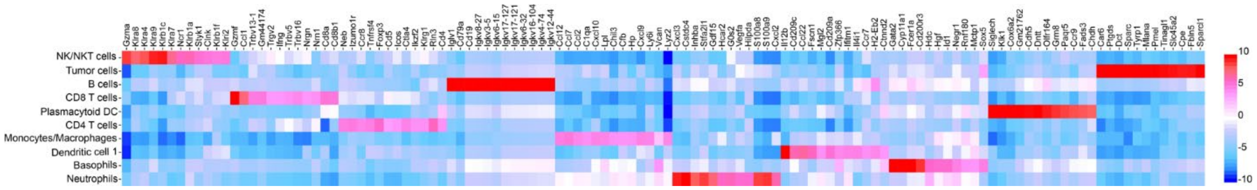


Figure S11. Heatmap of top 10 upregulated genes for each immune cluster.

| Antibody | Fluorophore | Company |
|---|---|---|
| CD3 | Alexa Fluor 488 | BioLegend (CA, USA) |
| CD11b | Spark NIR 685 | BioLegend (CA, USA) |
| CD8a | APC-Fire 750 | BioLegend (CA, USA) |
| CD4 | Brilliant Violet 785 | BioLegend (CA, USA) |
| CD25 | PE-Cy7 | BioLegend (CA, USA) |
| PD-1 | PE-Dazzle 594 | BioLegend (CA, USA) |
| CTLA-4 | Brilliant Violet 421 | BioLegend (CA, USA) |
| Granzyme B | APC | BioLegend (CA, USA) |
| CD45 | Pacific Blue | BioLegend (CA, USA) |
| Fixable Live/Dead | Zombie Aqua | BioLegend (CA, USA) |
| CD44 | Brilliant Violet 570 | BioLegend (CA, USA) |
| CD62L | Brilliant Violet 650 | BioLegend (CA, USA) |
| TRP2 | PE | Santa Cruz Biotechnology (TX, USA) |
| IFN-gamma | PerCP-Cy5.5 | ThermoFisher (IL, USA) |

Table S1. List of antibodies and fluorophores used for flow cytometry studies.
